# Nanoalgosomes from *Tetraselmis chuii*: Microalgal Extracellular Vesicles for UV Protection, Anti-Aging, and Skin Depigmentation

**DOI:** 10.1101/2025.07.25.666736

**Authors:** Paola Gargano, Sabrina Picciotto, Angela Paterna, Samuele Raccosta, Estella Rao, Daniele Paolo Romancino, Giulia Smeraldi, Mauro Manno, Monica Salamone, Natasa Zarovni, Giorgia Adamo, Antonella Bongiovanni

## Abstract

The search for effective dermocosmetic treatments has recently been accompanied by the growing demand for ingredients that are both naturally derived and sustainable. In this context, microalgae have emerged as a promising biofactory, producing bioactive compounds for skin health, widely recognized for their antioxidant, anti-inflammatory, and anti-aging activities. To leverage these properties, besides the conventional use of microalgal mass or extracts, the isolation and application of their secretome, including the extracellular vesicles named “nanoalgosomes”, emerged as a novel and increasingly studied approach. EVs are membranous nanoparticles released by all cells and naturally efficient in the transport of both endogenous and exogenous bioactive molecules. Their unique biological properties make them ideal candidates for therapeutic and cosmetic applications. Here, we propose nanoalgosomes, as a sustainable and effective solution for innovative dermocosmetic treatments. In this study we exemplify the use of the microalgae *Tetraselmis chuii,* an edible, green and renewable bio-source of extracellular vesicles. We demonstrated the nanoalgosomes’ skin-health promoting potential, by employing the human skin cells as model to evidence the nanoalgosome ability to protect the cells from ultraviolet B (UVB) radiation-related damages, by reducing both the oxidative stress, and senescence-associated phenotype in UVB-exposed skin cells. Furthermore, nanoalgosomes modulate melanogenesis in UVB-stimulated melanocytes by downregulating tyrosinase, significantly reducing melanin content, supporting their role as photoprotective and pigmentation-modulating agents. Altogether, our results provide a solid foundation for the development of nanoalgosome-based formulations in dermocosmetic applications aimed at UV protection, anti-aging, and depigmentation, supporting the transition toward natural and sustainable skincare solutions.

## 1.0 Introduction

The dermocosmetic industry has been significantly influenced by the growing demand for eco-friendly products, driving a shift toward the use of natural and sustainable ingredients. While natural compounds are often perceived as safer and more effective for skin health, their large-scale use presents significant environmental challenges, including extensive water consumption, land exploitation, and biodiversity depletion. Additionally, natural extracts often suffer from variability in composition due to differences in cultivation methods, climate conditions, and extraction processes, raising concerns about their standardization.

To overcome these limitations, innovative bio-based approaches from cellular biotechnology are bridging sustainability and performance. One of the most promising approaches in this field is the use of extracellular vesicles (EVs) as innovative carriers for bioactive molecules in both therapeutic and cosmetic applications. EVs are nanosized lipid-bilayer delimited vesicles physiologically released by all cell types, playing a crucial role in cell-to-cell communication by transporting all kinds of biomolecules, at most represented by proteins, lipids, and nucleic acids [1,2]. In recent years, EV-based formulations emerged as alternative to traditional synthetic lipid-based systems, such as liposomes and lipid nanoparticles [3–5]. While liposomes have been widely used in cosmetic formulations due to their ability to encapsulate and deliver active ingredients, they lack the intrinsic biological functionality of EVs [6]. Additionally, EVs have been shown to improve stability and protection for encapsulated compounds, making them highly attractive for the management of skin-related disorders and dysfunctions. Despite these advantages, mammalian-derived EVs face challenges related to large-scale production, standardization and safety, and are currently prohibited in Europe and China for the cosmetic application. To find more sustainable and scalable EV sources, researchers have turned to alternatives, one of which is remarkably presented by microalgae. Microalgae are photosynthetic microorganisms, rich in valuable metabolites, including antioxidants, polysaccharides, peptides, and pigments, some of which have been shown to exert beneficial effects for skin health. For that reason, microalgae have gained increasing attention as a sustainable and versatile source of bioactive compounds for cosmetics. Nanoalgosomes (ALG) are microalgae-derived extracellular vesicles that share fundamental properties with EVs (including exosomes), such as nanoscale size, membrane-bound structure, specific EV biomarkers, and the ability to be internalized by various human cells. Importantly, ALG shown to be highly stable in biological fluids, an additional feature to confirm their promising potential for applications in the cosmetic and pharmaceutic industry [7,8]. In this study, we investigated the potential of *Tetraselmis chuii*-derived nanoalgosomes (*TC*-ALG) as an innovative, biocompatible, sustainable and bio-active nanocarriers for skin health. The premises of the current study were built upon our previous studies that demonstrated that *TC*-ALG exhibit pronounced antioxidant and anti-inflammatory properties, as well as an *in vivo* biocompatibility.[7,9,10]. Human skin is an organ that has many roles like thermoregulation and regulation of hydration, while it especially works as a barrier to protect from external factors such as mechanical stress, infection and UV radiations [11,12]. The natural defence against UV rays is exerted by a specific type of cells situated in the epidermis, the melanocytes. When stimulated by UV rays, they produce melanin, which is then transferred to keratinocytes and works like a natural sunscreen [13,14]. There are three different types of UV rays: UVC, that are the most dangerous, typically shielded by ozone; UVA (320-400 nm) that can reach the dermis and cause redness, furrows, aging and melanoma; and UVB (280-320 nm) that affect the epidermis, determining tan, but can also cause cell damage and tumours [13,15–17].

Prolonged exposure to UV rays results in the rise of intracellular reactive oxygen species (ROS) and the activation of inflammatory and apoptotic pathways [18,19]. These molecular events promote the accumulation of senescent cells, characterized by increased senescence-associated (SA)-β-galactosidase activity and secretion of pro-inflammatory mediators [20]. From a more strictly dermocosmetic perspective these events can also lead to age-related skin issues such as dark spots, wrinkles, and loss of elasticity. Overall, all these events are responsible for the phenomenon known as photoaging [21,22].

In this context, recent evidence demonstrated that EVs (*e.g.* exosomes) derived from human stem cells can counteract senescence in fibroblasts, reduce matrix metalloproteinases expression, enhance collagen synthesis, and restore extracellular matrix homeostasis [20]. Among the various active ingredients, marine microalgae chemical extract has also been shown to possess photoprotective activity, improving cell viability, rebalancing the oxidation state and reducing the melanin content after UVB rays’ exposure [23,24]. In this study, we showed that, similarly to the extract effect, but without the use of solvents or depletion of the microalgal biomass, *TC*-ALG improves human skin cell viability after UVB exposure while exerting potent antioxidant effects. We also found that *TC*-ALG possess anti-melanogenic properties, likely modulating the expression of tyrosinase, a key enzyme in melanin biosynthesis, with a depigmenting effect that is comparable to that of arbutin, a widely used cosmetic ingredient. These findings, along with the advantages of a controllable, efficient, and renewable bioprocess, position nanoalgosomes as a next-generation, sustainable alternative for advanced dermocosmetic treatments against photoaging.

### 2.0 Materials and methods

### 2.1 *TC*-ALG isolation

*TC*-ALG isolation was performed from a 6L culture of the microalga *Tetraselmis chuii* (CCAP 66/21b), grown in F/2 medium, after 30 days of cultivation at a temperature of 20°C ± 2°C with continuous air flow and exposed to white light with a 14h light and 10h dark period [25]. The bottles were gently agitated every day to homogenize the cultures and 0.22 μm filters were placed at the bottle inputs to keep microalgae cultures in axenic conditions. Cell growth was evaluated every week using OD600 and cell viability using fluorescein diacetate (Thermo Fisher). Extracellular vesicle isolation and concentration were performed using the Repligen KrosFlo® KR2i TFF System (Spectrum Labs, Los Angeles, CA, USA) with three different GE Healthcare polysulfone hollow fiber membranes, as described by Adamo et al. (2021) [7,25]. The obtained samples were diafiltrated seven times to make a change buffer in phosphate-buffered saline without calcium and magnesium (PBS).

### 2.2 *TC*-ALG characterization

*TC*-ALG were characterized with biophysical and biochemical methods, following MISEV2018 and 2023 guidelines [4,26].

*Nanoparticle tracking analysis* (NTA) was performed by NanoSight NS300 (Malvern Panalytical, UK) to assess *TC*-ALG concentration and the size distribution. *TC*-ALG were diluted in HPLC-grade water (Sigma Aldrich), followed by 0.22 µm filtration to ensure the absence of particles (Whatman Anotop filters), to have 20-120 particles per frame. Five 60-second videos were analysed for each sample (n = 3) by NTA software Build 3.1.46, in light scattering, using a detection threshold set to 5 and the camera level at 15-16.

*Nanoparticle tracking analysis in fluorescence* (F-NTA) was determined using Di-8-ANEPPS-labelled *TC*-ALG (Ex/Em ∼465/635 nm, Thermo Fisher Scientific), a dye that emits fluorescence when in presence of EVs lipidic membranes, following Adamo et al. protocol [7]. In brief, *TC*-ALG were incubated 1h at room temperature with Di-8-ANEPPS and analysed by NTA equipped with a 500LP filter (480 nm laser).

*BCA assay* was used to evaluate the protein concentration of *TC*-ALG preparations using micro-bicinchoninic BCA Protein Assay Kit (Thermo Fisher Scientific). This is a colorimetric assay which uses a bovine serum albumin as standard curve. The BCA reaction solution absorbance was measured by GloMax Discover Microplate Reader at 560 nm.

*DetectEV assay* was used to evaluate the bioactivity and membrane integrity of three different batches of *TC*-ALG. 2 × 10¹⁰ nanoalgosomes were incubated with 18 µM fluorescein diacetate (FDA; Sigma-Aldrich) in 0.2 µm-filtered phosphate-buffered saline (PBS) without Ca²⁺ and Mg²⁺, in a final volume of 200 µL in a 96-well microplate. Fluorescence emission was monitored for up to 3 hours using the blue filter (excitation: 488 nm) of the GloMax Discover Microplate Reader (Promega). *TC*-ALG enzymatic activity was determined according to the protocol described by Adamo et al., 2025 [25,27]. A standard calibration curve was established using fluorescein sodium salt (Ex/Em ∼490/514 nm; Sigma-Aldrich), diluted in Ca²⁺- and Mg²⁺-free PBS to final concentrations ranging from 300 nM to 0 nM, with a total volume of 200 µL. Fluorescence was measured under the same conditions as the assay. Extracellular vesicle-associated enzymatic activity was expressed in enzymatic units (U), defined as the amount of fluorescein generated per minute (nM/min). This value was calculated by interpolating the fluorescence intensity measured at the end of the incubation period onto the standard curve and dividing the resulting fluorescein concentration by the total incubation time (180 minutes).

*Atomic Force Microscopy* (AFM) to assess *TC*-ALG morphology was performed using Nanowizard III scanning probe microscopy (JPK instruments) with a 15 µm z-range scanner. *TC*-ALG were resuspended in MilliQ water to obtain a specific concentration of 5×10^11^ particles/mL and 30 μL aliquot of the sample was deposited for 10 minutes onto freshly cleaved mica. The surface was then gently dried under a nitrogen stream. Images were acquired using an NSC-15 cantilever (Mikromasch), featuring a typical spring constant of 40 N/m and a tip radius of 8 nm.

*Immunoblotting* was performed with protein extracts from *TC*-ALG and microalgal cell lysates and separated by sodium dodecyl-sulfate polyacrylamide gel electrophoresis (SDS– PAGE). Prior to electrophoresis, 5 μg of protein from each sample were combined with 5x loading buffer (containing 0.25 M Tris-HCl pH 6.8, 10% SDS, 50% glycerol, 0.25 M dithiothreitol, and 0.25% bromophenol blue). The mixtures were then heated to 100 °C for 5 minutes and then loaded in the SDS-PAGE. Electrophoretic separation was followed by proteins transfer onto polyvinylidenfluoro (PVDF) membranes. Membranes were blocked at room temperature for 1h in a solution of 3% bovine serum albumin (BSA) in TBS-T (Tris-buffered saline with 0.05% Tween-20). Primary antibodies against H⁺-ATPase (1:1000, Agrisera), Alix (clone 3A9, 1:150, Santa Cruz), and β-actin (clone 10-B3, 1:500, Sigma-Aldrich), all diluted in blocking solution, were incubated overnight at 4°C and 2h at room temperature. After washing steps, membranes were incubated for 1h with species-specific HRP-conjugated secondary antibodies (Cell Signaling). Blots were then washed thoroughly (four washes for 20 minutes, in TBS-T), and signal detection was carried out using the SuperSignal Pierce ECL (Thermo Fisher Scientific).

### 2.3 *Tetraselmis chuii* extract preparation

*Tetraselmis chuii* cultures were also used to prepare a methanolic extract. Starting from 60 mg of dry biomass, 10 mL of methanol were added, and the sample was subjected to magnetic stirring for 30 minutes. After agitation, the sample was centrifuged at 3000 × g for 5 minutes. The resulting supernatant was then carefully removed, and the extraction steps were repeated until a clear methanolic solution was obtained. The collected methanolic fractions were pooled and concentrated using a rotary evaporator (IKA RV 10, Germany) to eliminate the solvent under reduced pressure. After evaporation, the resulting residues were subjected to overnight vacuum drying to ensure complete removal of any residual solvent. A total of 4 mg (dry weight) of the methanolic extract was then resuspended in a water and methanol mixture (2:1, v/v) and an equal volume of diethyl ether was then added. Following vortexing, the mixture was left to enable phase separation. The upper organic phase was collected, and anhydrous sodium sulphate was added to remove residual water. Final solvent removal was carried out using a rotary evaporator until complete evaporation was achieved. After 24 hours, the dry extract was dissolved in DMSO (vehicle) at 400 µg/ml (dry dry weight *TC*-extract in DMSO) and stored at –20°C for further analysis.

### 2.4 Cell culture

For this study, normal human dermal fibroblasts (NHDF) and human melanoma cells (A375) were chosen as cellular models. The NHDF and A375 cell lines were maintained in culture at 37°C with 5% CO_2_ in Dulbecco’s Modified Eagle’s Medium (DMEM) (Sigma-Aldrich), containing 10% (v/v) fetal bovine serum (FBS) (Gibco, Life Technologies), with 2 mM L-glutamine, 100 U/ml penicillin, and 100 mg/ml streptomycin (Sigma-Aldrich).

### 2.5 Viability of human skin cells *in vitro* following *TC*-ALG treatment

The NHDF and A375 cells were plated in 96-well plates at a density of 2×10^3^ cells/well and maintained in culture for 24h in their own medium at 37°C with 5% CO_2_. The viability assay was conducted by incubating *TC*-ALG at two different concentrations, 0.5 and 2 μg/mL (corresponding to about 10^9^ and 4×10^9^ *TC*-ALG/mL) [7,28]. After 24h of culture, the treatment was performed with the two concentrations of *TC*-ALG and incubated for 24, 48 and 72 hours. A group of cells had their medium replaced with fresh medium and were used as a negative control (100% viability). Cell viability was evaluated using CellTiter 96 Aqueus One Solution Reagent (Promega) and after 3h of incubation the absorbance (OD, Absorption at 490 nm) was calculated. The optical density was used to calculate the percentage of live cells using the following formula:

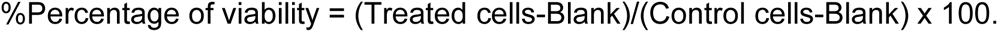

The experiment was performed in triplicate and with three different *TC*-ALG’ batches and ± SD of the three replicates was calculated.

### 2.6 Cellular uptake of *TC*-ALG and endosomal localization study

The cells were plated in 24-well plates containing sterile cover slips at a density of 3×10^4^ cells/well and incubated for 24 hours at 37°C with 5% CO_2_. After 24 hours, *TC*-ALG labelled with Di-8-ANEPPS were incubated at concentrations of 0.5 and 2 μg/mL [9]. After 3, 6 and 24 hours, two washes with PBS were performed, followed by fixation with 3.7% paraformaldehyde for 15 minutes and then two more washes with PBS. The nuclei were stained in blue with DAPI (VECTASHIELD Mounting Medium, Promega) [7]. The localization of *TC*-ALG was evaluated using fluorescence microscopy (EVident Olympus).

To evaluate the *TC*-ALG colocalization with the protein CD63, an endosomal marker, first *TC*-ALG were labelled with PKH26 (Ex/Em ∼551/567 nm, Sigma Aldrich), as described in Adamo at al 2021 and Picciotto et al 2022 [7,29]. Briefly, PHK26 was prepared at a final concentration of 3 μM in Diluent C (supplied by Sigma-Aldrich with the PKH26 kit). This labelling solution was then incubated with nanoalgosome samples at a concentration of 5×10¹⁰ particles/mL for 1 hour at 37 °C. Cells were grown at a density of 3×10^4^ cells/well in 24-well plates containing sterile coverslips in complete medium and, after 24h, cells were incubated with 2 μg/mL of PKH-26-labeled *TC*-ALG at 37°C for 24h [7,29]. Cells were fixed with 3.7% paraformaldehyde for 15 min at room temperature (RT) and washed twice with PBS. Cells were then permeabilized with 1% Triton X-100 in PBS for 10 min at RT. Afterwards, cells were incubated with blocking solution (1% bovine serum albumin in PBS) for 30 min at room temperature. Cells were incubated in a primary antibody (anti-CD63, clone MX-49.129.5), After three washes with PBS, cells were then incubated with a secondary antibody (AlexaFluor-488; Thermo Fischer Scientific) and diluted in blocking solution (1:50) for 2h at room temperature. Coverslips were mounted with a drop of Vectashield Mounting Medium with DAPI (Sigma-Aldrich). *TC*-ALG intracellular localization was monitored by fluorescent microscopy (EVident Olympus) [7].

### 2.7 UVB irradiation

UVB exposure was performed using a UVB lamp (312 nm peak) at 300 J/m^2^ for 42 seconds, setting the exposure to have about 40% of cell death. The cells (*i.e.*, NHDF and A375 cells) were kept in a small volume of phosphate-buffered saline (PBS) during irradiation [30,31]. After UVB exposure, PBS was replaced with fresh medium, and cells were kept in culture for 24 and/or 48h.

### 2.8 Assessment of antioxidant activity of *TC*-ALG in UVB-irradiated skin cells

Intracellular ROS levels of living cells were determined using 2,7-dichlorofluorescein diacetate (DCFH-DA; Sigma-Aldrich). This non-fluorescent probe emits fluorescence in the presence of ROS in a directly proportional manner, allowing a quantitative assessment of oxidative stress by ROS evaluation. The NHDF and A375 cells were plated in a 96-well plate at a density of 5×10^3^ cells/well and incubated at 37°C with 5% CO_2_ in William’s E medium without phenol red (Gibco™). After 24h, both cell lines were treated with two different concentrations of *TC*-ALG (0.5 and 2 μg/mL). As negative control, untreated cells were incubated with fresh medium to set ROS basal level, and as antioxidant-positive control, other cells were treated with 70 µg/mL of arbutin, due to its proven antioxidant effect. After 24h of treatment, cells were exposed to UVB light, as described before. Next, cells were incubated with DCFH-DA, at a concentration of 40 μM diluted in medium without FBS for 1h at 37°C with 5% CO_2_. Moreover, cells treated with a 250 μM solution of tert-butyl hydroperoxide (TBH, Sigma-Aldrich) diluted in medium without FBS and incubated for 1h at 37°C with 5% CO₂, were used as oxidative-positive control using an oxidizing agent distinct from UVB radiation. After 24h, both cell lines were treated with two different concentrations of *TC*-ALG (0.5 and 2 μg/mL) and with three different concentrations of *TC*-extract (1, 2, and 5 μg/mL of dry weight of *TC*-extract in DMSO, in NHDF only). After 1h, intensive washes with PBS with calcium and magnesium were performed and fluorescence was read (Ex/Em ∼485/538 nm) using the GloMax Discover Microplate Reader [9,25]. The relative percentage of intracellular ROS was normalized with respect to untreated cells (negative control).

### 2.9 Photoprotective role of *TC*-ALG on cell viability after UVB exposure

The NHDF and A375 cells were seeded in 96-well plates at a density of 5×10^3^ cells/well and maintained in culture for 24h in their own medium at 37°C with 5% CO_2_. Two concentrations of *TC*-ALG (0.5 and 2 μg/mL) were used for the experiment, with treatments carried out using three different strategies: 24h of pre-incubation with *TC*-ALG followed by UVB exposure, *TC*-ALG incubation during the UV exposure and the third *TC*-ALG incubation after UVB exposure. As negative control, untreated cells were incubated with fresh medium (100% viability); as positive control cells were treated with 70 µg/mL (257 µM) of arbutin (Sigma Aldrich) [32]. After UVB exposure, cells were kept in culture for 24h for the three conditions, next, the photoprotective role of *TC*-ALG was evaluated using CellTiter 96 Aqueus One Solution Reagent, as described before [24].

### 2.10 Anti-photoaging effect of *TC*-ALG on UVB-induced senescent human dermal fibroblasts

Senescence-associated β-galactosidase (SA-β-gal) staining was carried out using a commercially available kit (Sigma-Aldrich CS0030) based on a histochemical stain for β-galactosidase activity at pH 6 using its substrate X-gal (5-bromo-4-chloro-3-indolyl-beta-D-galacto-pyranoside). Following the manufacturer’s protocol, cells were seeded in 24 well/plate at a density of 1×10^4^ cell/well, and after 24h cells were treated with/without 0.5 µg/mL of *TC*-ALG; this concentration was chosen following antioxidant and photoprotective results. After 24h cells were exposed to UVB rays (300 J/m^2^) and leaved in culture for 24h. To perform the SA-β-gal staining, cells were first washed twice in PBS and subsequently fixed with Fixation Buffer for 6 minutes at room temperature. After fixation, cells were washed three times with PBS, stained with 1 mL of Staining Mixture and incubate for 24h at 37 °C in a non-CO₂ humidified incubator. Stained cells were then visualized under a light microscope (OLYMPUS, CKX3-SLP), and SA-β-gal-positive cells were quantified acquiring 14 fields/condition (≈800 nuclei/condition) and analysing by ImageJ software (National Institutes of Health, USA). The percentage of SA-β-gal-positive cells (senescent cells) was calculated relative to the total number of cells counted [33,34].

### 2.11 Anti-melanogenic effect of *TC*-ALG in human melanoma cells

The A375 cell lines were plated in 6-well plates at a density of 1×10^5^ cell/well and incubated at 37°C with 5% CO_2_ in complete DMEM reaching 60% of confluency. Next, cells were pre-treated with 0.5 and 2 μg/mL of *TC*-ALG, and with 70 µg/mL of arbutin. As negative controls, untreated cells were used to set the melanin basal level. After 24h, cells were exposed to UVB rays or with 200 nM of α-melanocyte stimulating hormone (α-MSH, Sigma Aldrich), to stimulate melanin production. After 48h, melanin extraction was performed following Ruri Lee et al [32]. Briefly, cells were detached using a cell scraper, counted using Thoma counting chamber, and 1 million cells per condition were collected and centrifuged at 1000 rpm for 5 minutes. The resulting pellets were dissolved in 1 N NaOH containing 1% DMSO. Each sample were then incubated at 80°C with shaking at 250 rpm for 1h. Cell lysates were transferred to a 96-well plate, and melanin content was measured by reading absorbance at 405 nm using a GloMax Discover Microplate Reader. A standard curve of melanin (0-100 µg/mL) was prepared using a synthetic melanin (Sigma Aldrich) dissolved in 1 N NaOH [32].

### 2.12 Anti-tyrosinase activity of *TC*-ALG in human melanoma cells

Western blot analysis: for tyrosinase expression study, A375 cells seeded and treated using the same experimental setting described in section 2.9. After 24h, cells were lysed, and total proteins were extracted by the RIPA buffer and separated by 10% SDS-PAGE. A total of 10 μg of cell lysate were mixed with 5X loading buffer (0.25 M Tris-Cl pH 6.8, 10% SDS, 50% glycerol, 0.25 M dithiothreitol (DTT), 0.25% bromophenol blue) and heated at 100°C for 5 minutes. Proteins were electro-transferred onto a polyvinylidene difluoride membrane (PVDF), and the non-specific antibodies were blocked by incubating with Odyssey Blocking Buffer (Li-COR) in TBST 1X (50 mM Tris HCl pH 8.0, 150 mM NaCl with 0.05% Tween-20). Anti-Tyrosinase (TYR) (Sigma Aldrich, ab61294) primary antibody was incubated 1:1000 at 4°C over night and 2h at room temperature (RT). Afterwards, 1:10000 IRDye® 800CW Goat anti-Mouse IgG secondary antibody (LI-COR) was incubated for 1h at RT [32]. The immunoblot was revealed with Odyssey LI-COR and the densitometric analysis was conducted using Image Lab 6.1 BIO-RAD software, in which tyrosinase band density was normalized to the total protein levels determined by Ponceau S staining.

For the immunofluorescence study, A375 were seeded in 24-well plate with sterile coverslips, at a density of 3×10^4^ cell/well and incubated at 37°C with 5% CO_2_ in complete DMEM. Cells were treated with 0.5 μg/mL of *TC*-ALG and after 24h of incubation, 200 nM of α-MSH was added for 24h. Next, cells were fixed with 3.7% paraformaldehyde for 15 minutes and then were washed twice with PBS. Cells were permeabilized with 0.1% Triton X-100 (Sigma Aldrich) in PBS for 5 minutes at RT and then blocked with 1% BSA for 30 minutes at RT. Anti-tyrosinase antibody was added o/n at 4°C diluted 1:1000 in 1% BSA [35]. Cells were then incubated with a secondary antibody (AlexaFluor-488; Thermo Fischer Scientific) and diluted in blocking solution (1:50) for 2h at room temperature. Coverslips were mounted with a drop of Vectashield Mounting Medium with DAPI (Sigma-Aldrich) .and images were taken with Evident Olympus fluorescence microscope. The fluorescence intensity analysis was performed using three different fields/condition, analysing a minimum of 18 nuclei per field, using ImageJ software.

### 2.13 Statistical analysis

All experiments were repeated independently at least 3 times, with data expressed as mean ± SD. Statistical analysis and graphs were conducted using GraphPad Prism 8 (GraphPad Software Inc., USA). Statistical comparisons were performed using one-way or two-way ANOVA (three or more groups), with Tukey’s or Dunnett’s multiple comparisons test. Statistical significance was considered at p<0.05. Significance levels were denoted as follows: *p<0.05, **p<0.01, ***p<0.001, and ****p<0.0001. Non-significant differences are indicated as "ns".

## 3.0 Results

### 3.1 Isolation and quality checking of *Tetraselmis chuii*-derived nanoalgosomes

The isolation of nanoalgosomes from the conditioned media of the marine chlorophyte microalgae *Tetraselmis chuii* was carried out using Tangential Flow Filtration (TFF), with sequential 650 nm and 200 nm pore-size cartridges, followed by diafiltration and concentration in phosphate-buffered saline (PBS) using a 500 kDa cut-off membrane [7,9,36]. These preparative steps allowed to concentrate the small EV fraction more than 1000 fold, by obtaining 5 mL samples containing small EVs in PBS from an initial culture volume of 6 L of *Tetraselmis chuii* conditioned media [7,9,26]. The comprehensive characterization of the obtained EV preparation was conducted following MISEV guidelines, allowing us to produce quality-controlled *TC*-ALG preparations in terms of protein and nanoparticle concentration, size distribution, and expression of EV biomarkers (**Fig. 1**), with high batch-to-batch reproducibility [26].

**Figure 1:**
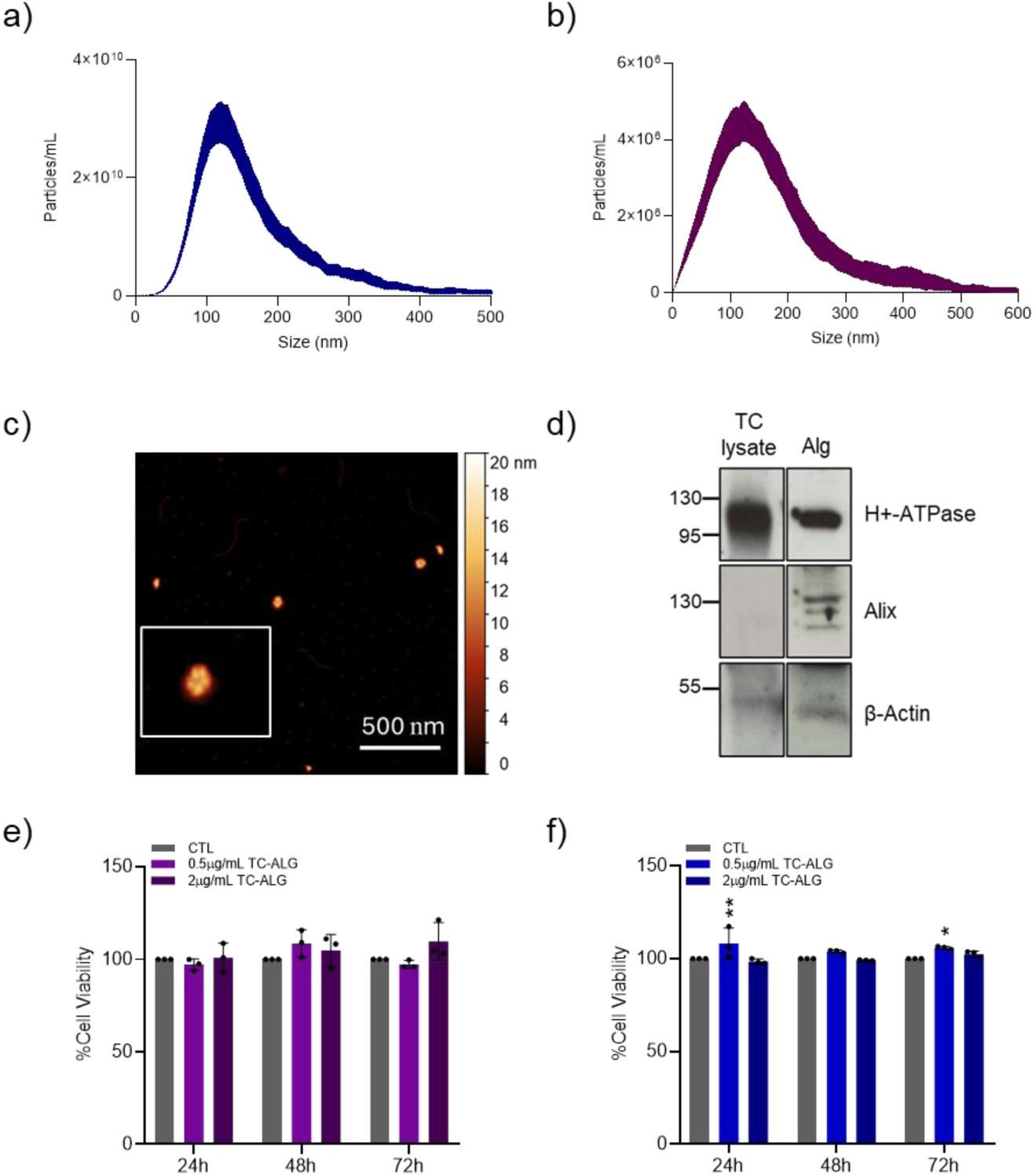
*TC*-ALG quality check and biocompatibility in vitro assay in skin cells. a) *TC*-ALG concentration and size distribution measured with NTA (blue deviation is relative to five measurements per sample); b) size distribution of Di-8-ANEPPS-labelled *TC*-ALG measured with Fluorescent (F)-NTA (purple deviation is relative to five measurements per sample); c) morphological evaluation of *TC*-ALG using AFM; d) *TC*-ALG marker expression (*e.g.*, H+-ATPase, Alix and β-Actin) analysed with immunoblot. *TC*-ALG biocompatibility evaluation in NHDF (e) and A375 (f) cell, by MTS assay. Statistical analyses were performed using two-way ANOVA (comparing control *vs*. treatments at each time point). Differences are reported as not significant (ns) or as *p < 0.05, **p < 0.01, where indicated. All the images are representative results of three independent replicates.

Across the *TC*-ALG preparations (n = 3), the protein content was 120 ± 20 µg/mL, the particles concentration 5×10^11^ ± 1×10^10^ particles/mL and the size distribution of 110 +/- 10 nm (mode), as measured by Nanoparticles Tracking Analysis (NTA) in scattering mode (**Fig. 1a**). NTA also indicated that the vesicles displayed a narrow size distribution, consistent with a uniform population of small EVs. Fluorescence NTA analysis was performed using Di-8-ANEPPS-labeled *TC*-ALG, confirmed the presence of membrane-bound particles within the same size range, validating their identity as EVs (**Fig. 1b**). Furthermore, the ratio between EV particle number and total protein content was consistent across all *TC*-ALG batches (n = 3), with 1 µg of EV protein corresponding to 5–10 × 10⁹ particles. This falls within the optimal range for small EVs, as proposed by Sverdlov [37]. These results were further validated through atomic force microscopy (AFM), which confirmed the *TC*-ALG morphology as typical of small EVs (**Fig. 1c**). Furthermore, protein profiling through SDS-PAGE and immunoblotting detected canonical EV markers such as H⁺-ATPase and Alix as well as β-Actin. These molecular signatures support the successful isolation of highly enriched EV preparations (**Fig. 1d**). Collectively, these results confirm that the nanoalgosomes derived from *Tetraselmis chuii* possess characteristic nanoscale dimensions, structural integrity, and marker protein expression typical of small EVs. Moreover, *TC*-ALG were additionally characterized using the functional DetectEV assay, which enables the assessment of both bioactivity and membrane integrity among different nanoalgosomes preparations, yielding an esterase-like activity equal to 1.5 ± 0.3 nM/min [27].

### 3.2 *TC*-ALG biocompatibility and internalization in human skin cells

The biocompatibility of *TC*-ALG was evaluated in normal human dermal fibroblasts (NHDF) and human melanoma cells (A375) using the MTS (3-(4,5-dimethylthiazol-2-yl)-5-(3-carboxymethoxyphenyl)-2-(4-sulfophenyl)-2H-tetrazolium) assay. Once confirmed the non-cytotoxic nature of *TC*-ALG in these skin cells (**Fig. 1e and f**), we performed an uptake study using a green-fluorescent version of *TC*-ALG labelled with the lipophilic dye Di-8-ANEPPS. Specifically, we incubated NHDF and A375 cell lines with Di-8-ANEPPS-labelled *TC*-ALG using different concentrations (0.5 and 2 μg/mL) for various incubation periods (6 and 24 hours). Both human skin cells take up *TC*-ALG in a time- and dose-dependent manner (**Fig. 2a and b**). Indeed, the green fluorescence relative to *TC*-ALG become evident after 6 hours and is even more pronounced at 24h. Furthermore, we have observed the intracellular localization (**Fig. 2c**) of PKH26-labelled *TC*-ALG (in red) within the CD63-positive endosomal compartment, through immunostaining for the CD63 protein (in green) in line with previous findings by Adamo et al [7,9].

**Figure 2:**
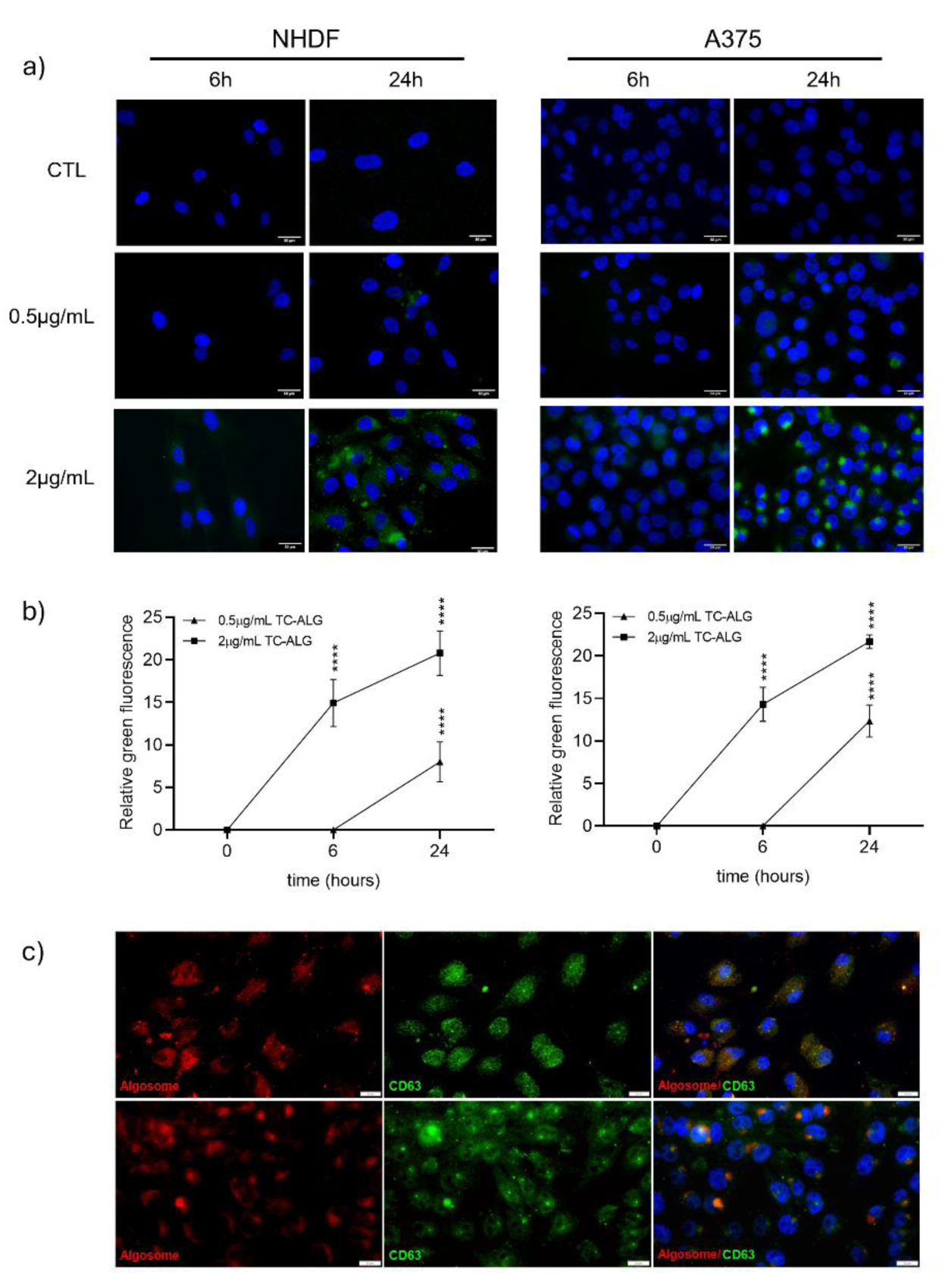
Cellular uptake and colocalization of CD63 and *TC*-ALG in human skin cells. a) Representative fluorescence microscopy images of NHDF and A375 cell lines (nuclei in blue), incubated with different concentrations of green labelled *TC*-ALG for 6 and 24h, at 37°C. b) Graphs of the quantification of the green fluorescence intensity relative to *TC*-ALG inside cells. The statistical analysis was conducted by two-way ANOVA using Dunnett’s multiple comparison test (control *vs*. *TC*-ALG) for each concentration and time; ****p <0.0001. c) Representative immunofluorescence microscopy images that show intracellular uptake of PKH26 fluorescent *TC*-ALG (in red) in NHDF and A375 cell lines (nuclei in blue) and the co-localization with the endosomal marker CD63 (in green). Scale bar 20 µm. All the images are representative results of three independent replicates.

### 3.3 *TC*-ALG contrasts the UVB-induced damage in skin cells

#### Antioxidant activity of TC-ALG

Several studies have explored the antioxidant potential of microalgal extracts in various biological systems [38]. Moreover, such microalgal extracts are already used in commercial cosmetic formulations; however, this raises sustainability challenges linked to the extraction and production processes required to obtain them.

As a preliminary step, we evaluated the antioxidant activity of methanolic extracts of our *Tetraselmis chuii* (*TC-*extract) to verify and confirm consistency with effects reported in the literature [39]. The *TC*-extract was tested at concentrations of 1, 2, and 5 μg/mL of dry weight in DMSO on NHDF cells; oxygen reactive species (ROS) levels were measured following oxidative stimulation with tert-butyl hydroperoxide (TBH). Consistently, the *TC*-extract exerted a significant antioxidant effect, reducing ROS production and better preserving cellular redox balance (**Supp. Fig. 1**). Having established the extract’s antioxidant potential as a control, we then focused our experiments on *Tetraselmis chuii*-derived extracellular vesicles (*TC*-ALG), which offer a solvent-free, cost-effective, and reproducible alternative to conventional microalgal extracts.

UVB rays are the ones responsible of DNA damage, solar erythema, photodamage, melanin synthesis and ROS production in cutaneous cells [40,41]. Previous studies demonstrated *TC*-ALG’ antioxidant activity *in vitro* (*e.g.* in breast cancer and non-tumorigenic mammary epithelial cell lines) and *in vivo* using H_2_O_2_ and TBH as oxidant agents, confirming also the absence of oxidant effect of *TC*-ALG per se [9]. In this study, the antioxidant role of nanoalgosomes was further investigated using UVB rays to induce ROS production in human skin cells. For this experiment, NHDF and A375 cells were pre-treated for 24h with two different concentrations of *TC*-ALG before UVB exposure. In parallel, cells were treated for 24h with arbutin, a well-established natural antioxidant widely used as a positive control in UVB-related effects studies. First, we confirm that arbutin treatment did not have any toxic effect in both cell lines used, performing an MTS assay and evaluating cell viability (**Supp. Fig. 2a and b**). Following the 24h of pre-incubation, cells were exposed to UVB radiation; UVB treatment significantly increased ROS production in both skin cell lines, similar to what observed upon the TBH-stimulation, confirming that UVB exposure severely affects cellular health by elevating intracellular ROS levels (**Fig. 3a and b**). Notably, 24h pretreatment with *TC*-ALG effectively reduces ROS levels in UVB-exposed cells, restoring their oxidative status to basal levels. Remarkably, nanoalgosomes exhibit an antioxidant effect comparable to that of arbutin (**Fig. 3a and b**). This finding suggests that nanoalgosomes may serve as a potential alternative to conventional antioxidant compounds in preventing oxidative stress.

**Figure 3:**
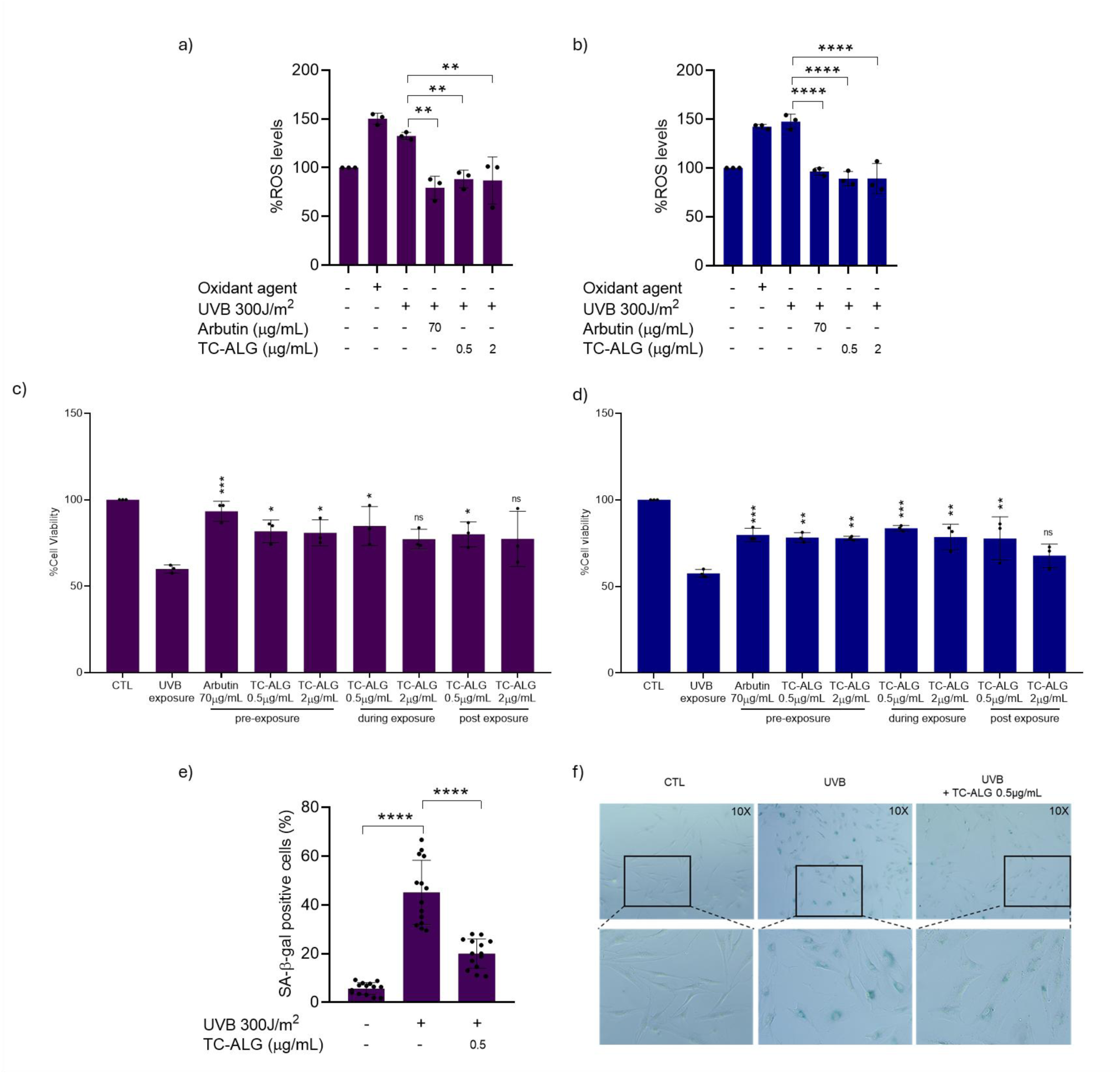
Antioxidant, photoprotective and anti-photoaging role of *TC*-ALG after UVB exposure in human skin cells. Panels (a) and (b) show the percentage of intracellular ROS level in NHDF and A375 cells, respectively, following treatment with *TC*-ALG (0.5 and 2 µg/mL), either alone or in combination with an oxidative stressor (250 µM TBH) or UVB exposure. In both cell lines, *TC*-ALG markedly reduced ROS accumulation, highlighting its antioxidant potential. Panels (c) and (d) illustrate the positive effect of *TC*-ALG (0.5 and 2 µg/mL) on cell viability before, during and after 24 hours of UVB irradiation in NHDF and A375 cells, respectively, indicating a clear photoprotective role. Panel (e) depicts the anti-photoaging activity of *TC*-ALG, expressed as the percentage of SA-β-gal-positive cells following UVB exposure, with or without *TC*-ALG treatment. Panel (f) presents representative bright-field images of SA-β-gal staining, visually confirming the reduction in senescence markers in *TC*-ALG-treated samples. The statistical analysis (UVB exposure *vs*. treatments) was conducted by ordinary one-way ANOVA *p < 0.05, **p < 0.01, ***p < 0.001, ****p > 0.0001. In e) data are presented as individual values, with each dot corresponding to the percentage of SA-β-gal positive cells per field (n = 14 fields per condition). Images are representative results of three independent replicates.

#### Photoprotective activity of TC-ALG

Given the previously observed antioxidant activity of *TC*-ALG in human skin cells, their potential to mitigate or prevent UVB-induced cellular damage was investigated by assessing cell viability in NHDF and A375 cell lines. Both cell lines were exposed to UVB radiation (300 J/m²) and subjected to three different *TC*-ALG treatment approaches to evaluate their potential effects against UVB-induced damages. Arbutin was included as a positive control [42]. **Figure 3c and d** show how UVB rays’ exposure resulted in a 45% reduction in viability for both skin cell lines compared to the not exposed control (CTL). Instead, treatments with *TC*-ALG, performed 24h before, during, or after UVB exposure, succeeded in significantly restoring viability levels. This result demonstrates the photoprotective role of *TC*-ALG, with an effect comparable to arbutin, even at a very low concentration (0.5 μg/mL, corresponding to 5×10⁹ particles/mL). The ability of *TC*-ALG to improve cell viability is likely linked to its capacity to modulate ROS production and mitigate their harmful effects. Notably, *TC*-ALG contrasts the UVB-induced cell damage when administered as a pre-treatment (before UVB exposure, preventing damage), as a co-treatment (during UVB exposure, contrasting ongoing damage), and as a post-treatment (after UVB exposure, restoring cell balance).

#### Anti-photoaging activity of TC-ALG

To investigate the potential of *TC*-ALG in protecting cell against UVB-related senescence (photoaging) we evaluated SA-β-galactosidase (SA-β-gal) activity in NHDF following UVB exposure (**Fig. 3e and f**). This assay exploits the elevated lysosomal β-galactosidase activity that characteristically accumulates in senescent cells and becomes detectable at pH 6.0. Cells were fixed and subsequently incubated with a chromogenic substrate, X-gal, which, upon enzymatic cleavage by SA-β-gal, produces an insoluble blue precipitate. The presence of blue-stained cells was then evaluated microscopically, and the percentage of SA-β-gal positive cells was quantified as relative to the total cell number. As expected, UVB rays considerably increased the number of SA-β-gal-positive cells, indicating enhanced early cellular senescence, while the treatment with 0.5 μg/mL *TC*-ALG significantly reduced SA-β-gal-positive cells compared to the UVB-only group, suggesting an anti-aging effect of these microalgae-derived EVs.

### 3.4 Anti-melanogenic and anti-tyrosinase effects of *TC*-ALG in human melanoma cells

To assess the anti-melanogenic role of *TC*-ALG, we treated human melanoma cells (A375) with two concentrations of *TC*-ALG and exposed them to UVB rays or alpha-melanocyte stimulating hormone (α-MSH) (**Supp. Fig. 2c**), a peptide released by the pituitary gland that bounds the melanocortin 1 receptor (MC1R), to stimulate melanin production. After 48 hours of combined treatment, we extracted melanin using the described protocol and measuring absorbance values at 405 nm. UVB treatment resulted in a 31% increase in melanin content, while treatment with arbutin (70 µg/mL) or *TC*-ALG (0.5 and 2 μg/mL) reduced melanin content respectively by 26.7%, 23.6% and 25.9% (**Fig. 4a**). α-MSH treatment induced an increase of melanin content of 42%, while arbutin and *TC*-ALG showed an anti-melanogenic effect of respectively of 29.6%, 25.35% and 30.9% (**Fig. 4b**). These results demonstrate the ability of *TC*-ALG to significantly reduce melanin content, and it is important to highlight that this effect is perfectly comparable to the arbutin, a depigmenting agent already used in cosmetic products that also have a moderate effect on modulating tyrosinase expression and a strong ability in inhibit tyrosinase activity.

**Figure 4:**
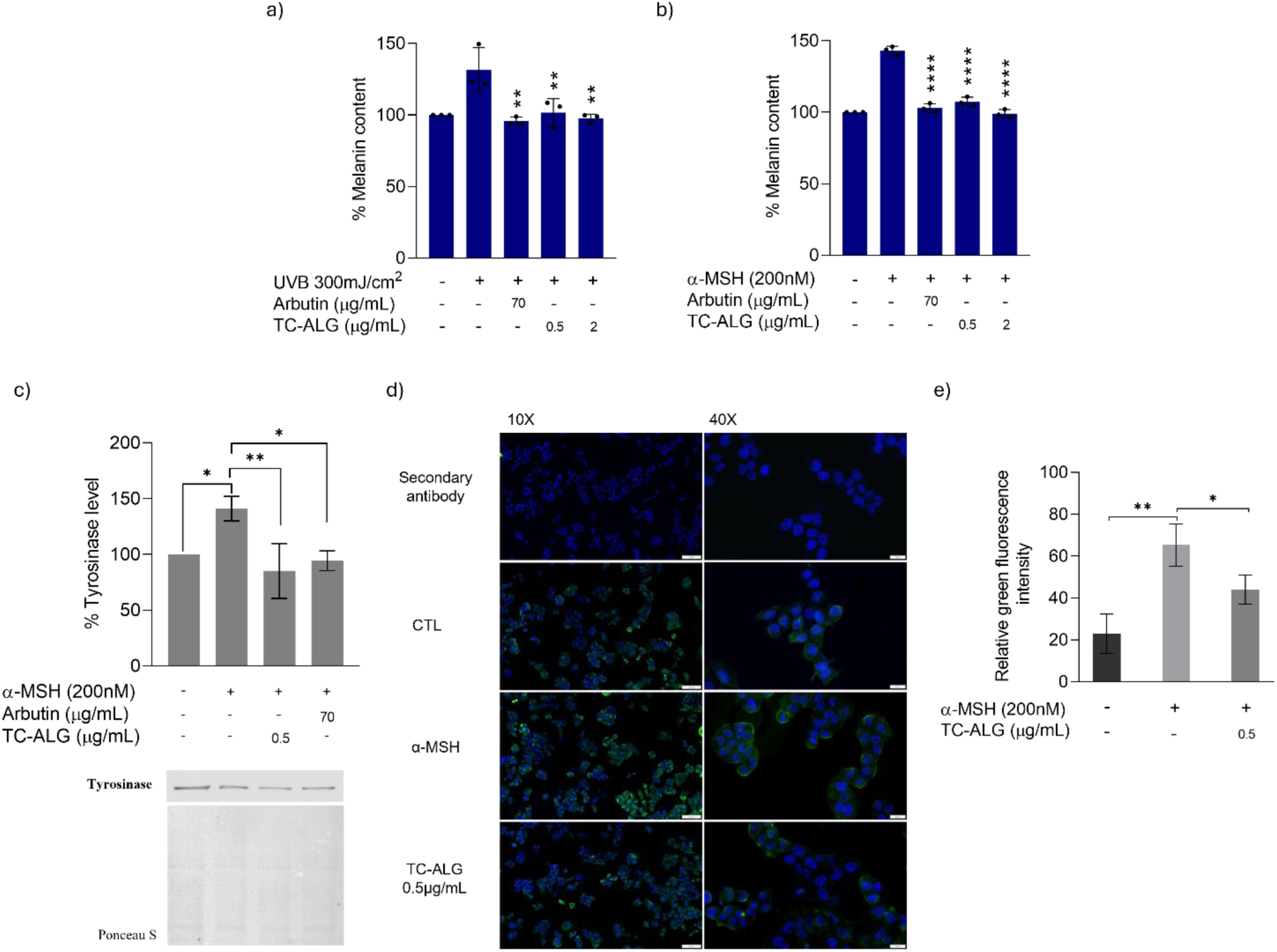
Anti-melanogenic effect of *TC*-ALG and tyrosinase expression in human melanoma cells after melanin stimulation through UVB or α-MSH. The graph shows the percentage of melanin content extracted from A375 cells after UVB exposure (a) or stimulated with α-MSH (b), demonstrating the anti-melanogenic effect of *TC*-ALG. (c) Western blot and densitometric analysis of tyrosinase expression (TYR) after α-MSH stimulation. (d) Representative image showing immunofluorescence obtained using a fluorescent tyrosinase antibody (green). (e) Fluorescence intensity analysis was performed using ImageJ software. The statistical analyses were conducted by Ordinary one-way ANOVA *p < 0.05, **p < 0.01, ***p < 0.001, ****p < 0.0001, performing Dunnett’s multiple comparisons test (UVB exposure or α-MSH *vs*. *TC*-ALG or arbutin). Images are representative results of three independent replicates.

To further investigate the specific mechanism by which *TC*-ALG exert their anti-melanogenic activity, tyrosinase (TYR) expression levels were analysed in A375 cells pre-treated with 0.5 μg/mL of nanoalgosomes (the lowest concentration at which significant effects on melanin reduction were observed) or 70 μg/mL of arbutin, followed by stimulation with α-MSH. Densitometric analysis of immunoblot results revealed that the treatment with α-MSH led to an increase in tyrosinase expression levels (**Fig. 4c**). However, pre-treatment with *TC*-ALG effectively counteracted this effect, restoring tyrosinase expression to basal levels and resulting in an approximate 30% reduction in protein levels (**Fig. 4c, Supp. Fig. 3**). These results were also validated by immunofluorescence (**Fig. 4d and e**). The representative images show that α-MSH stimulation results in an increased green fluorescence intensity, while *TC*-ALG pre-treatment can reduce this fluorescence and so tyrosinase levels of about 33%. These results provide further evidence that *TC*-ALG modulates melanogenesis acting on tyrosinase expression and reinforces their potential as depigmenting agents.

### 4.0 Discussion

Microalgae are known to produce a wide range of bioactive compounds with different biological activity, most of them related to pigments [43,44]. In our previous investigations, *Tetraselmis chuii* showed the presence of neoxanthin, violaxanthin, lutein, β-carotene and chlorophyll [8]. All these pigments, known for their antioxidant and photoprotective activity, are already used in cosmetic formulations, and some of them showed the ability to reverse aging and melanogenesis mechanisms [44–47]. Moreover, *Tetraselmis chuii* and other microalgal chemical extracts, as well as liposome-based skin delivery technology of natural active ingredients of microalgae, are widely studied and actually used in cosmetics [48–50]. It has been observed, indeed, that the encapsulation of entire microalgae extracts, as well as their single pigments (*e.g.* fucoxanthin or astaxanthin), within liposomes, enhances their thermal stability, absorption, and activity in various cell lines and *in vivo* [48,51]. Our previous studies confirmed that nanoalgosomes possess innate antioxidant and anti-inflammatory activities, due to their endogenous cargo, *in vitro* and *in vivo* [9,10]. Here we demonstrate that these microalgae-derived EVs have a significative activity on contrasting UVB-induced damage in skin cells, acting on the reduction of oxygen reactive species content, and consequently increasing cell viability and reverting photoaging mechanisms.

Moreover, recently main focus of cosmetic and dermatological research turned to the formulation of depigmenting products to treat skin with dark spots caused by UV exposure or pathological conditions like melasma. Unfortunately, most depigmenting agents have shown limitations in term of human safety and ecological footprint: hydroquinone was banned in 2001 due to cytotoxic effect of its metabolites, kojic acid was banned in Japan because of the reported allergic reactions, and the use of arbutin is restricted starting from 2023 [52–55]. The anti-melanogenic effect, which is the ability to reduce the melanin content produced by melanocytes, can be attributed to various mechanisms (e.g., degradation of the pigment, inhibition of the melanogenesis-stimulating hormone, inhibition of tyrosinase expression or activity) [56]. Such anti-melanogenic activities have been tackled over the years by treating melanocytes with various microalgae extracts. For example, it has been demonstrated that the microalga *Pavlova lutheri* can reduce melanin content in B16F10 tumour cells, and that the extract of the cyanobacterium *Phormidium persicinum* displayed an anti-melanogenic properties in human volunteers [57,58]. Given these findings, and considering the pigment profile of *T. chuii*, we aimed to evaluate the potential anti-melanogenic effect of nanoalgosomes by measuring melanin content in human melanoma cells stimulated with UVB radiation or α-melanocyte-stimulating hormone. Results showed that nanoalgosomes can significantly reduce the melanin content, confirming a depigmenting activity of these small-EVs. Importantly, the anti-melanogenic effect and the photoprotective role of *T. chuii*-derived nanoalgosomes, exerted already at a very low concentration, are perfectly comparable to the effect of arbutin, a well-known anti-melanogenic, antioxidant and photoprotective compound. These results suggest that, similarly to microalgae extracts, microalgae-derived EVs possess diversified beneficial properties and may offer a natural and sustainable alternative to synthetic compounds currently used for skin health treatments. With respect to whole extracts, the use of the microalgal-derived EVs offer several advantages: their biogenesis is a physiological and reproducible process, ensuring strong batch-to-batch consistency; nanoalgosomes’ production is solvent-free, rapid, sustainable and renewable, aligning with dermocosmetic market demands for eco-compatible and cost-effective ingredients. Furthermore, extracellular vesicles are naturally evolved to mediate intercellular communication by delivering bioactive molecules, which makes them able of inducing cellular responses and exerting specific biological effects. Unlike extracts, EVs can enrich and protect natural active compounds from microalgae, facilitate their delivery, and enhance their bioavailability, thus boosting efficacy. A limitation of this study is that the photoprotective and anti-melanogenic effects of nanoalgosomes were assessed exclusively *in vitro*, using human skin cell lines. While this is a widely accepted and informative first step to evaluate efficacy, it does not fully represent the structural and functional complexity of human skin. To strengthen the translational relevance of our findings, future studies should include advanced models such as organ-on-chip platforms or *in vivo* human studies, which would help to define nanoalgosomes activity in physiologic conditions.

Overall, our findings highlight the potential of nanoalgosomes (*i.e*., *TC*-ALG and other microalgae-derived EVs) as multifunctional dermocosmetic ingredients. Specifically, we propose their use both to counteract UVB-induced cellular damage by rebalancing ROS levels, and to reduce melanin synthesis as depigmenting agents for treating existing UVB-induced hyperpigmentation, such as melasma. These properties make them promising candidates for the treatment of a range of cutaneous alteration.

## Supporting information

The file Suppementary materials contains three additional figures

## Data availability

The data supporting this study are available in the repository for open data on Figshare (https://doi.org/10.6084/m9.figshare.29683559.v1).

## Declaration of Generative AI and AI-assisted technologies in the writing process

During the preparation of this work the author(s) used ChatGPT in order to improve language and readability. After using this tool, the author(s) reviewed and edited the content as needed and take(s) full responsibility for the content of the publication.

## Acknowledgements

The authors acknowledge Nicolas Touzet and Helen Herbert (ATU, Ireland) and María D. Mayán Santos (CINBIO – University of Vigo, Spain) for support and precious advice, and Luca Caruana (CNR-IRIB, Italy) for administrative support.

## CRediT

AB: Writing - Review & editing, Supervision, Methodology, Conceptualization. PG: Writing - original draft, Methodology, Investigation, Formal analysis, Conceptualization. SP: Writing - Review & editing, Methodology, Investigation, Formal analysis, Conceptualization. GA: Writing – Review & editing, Supervision, Investigation. MM: Writing - Review, Investigation. AP, SR, ER, DPR, GS, MS: Writing - review, Investigation. NZ: Writing - review & editing.

