## Supplementary material for "Nanoalgosomes from *Tetraselmis chuii*: Microalgal Extracellular Vesicles for UV Protection, Anti-Aging, and Skin Depigmentation": The file Suppementary materials contains three additional figures

**b Cell-Tech HUB and Institute of Biophysics (IBF) - National Research Council of Italy (CNR), Palermo 90146, Italy**

**c EVEBiofactory at Cell-Tech HUB, National Research Council of Italy (CNR), Palermo, Italy**

**1 These Authors contributed equally to this work and shared the first authorship**

#### Keywords

Microalgae-derived extracellular vesicles; Exosomes; Nanoalgosomes; Skin photoprotection; UVB-induced damage; Dermocosmetic formulations; Melanogenesis modulation.

#### Supplementary Figure 1

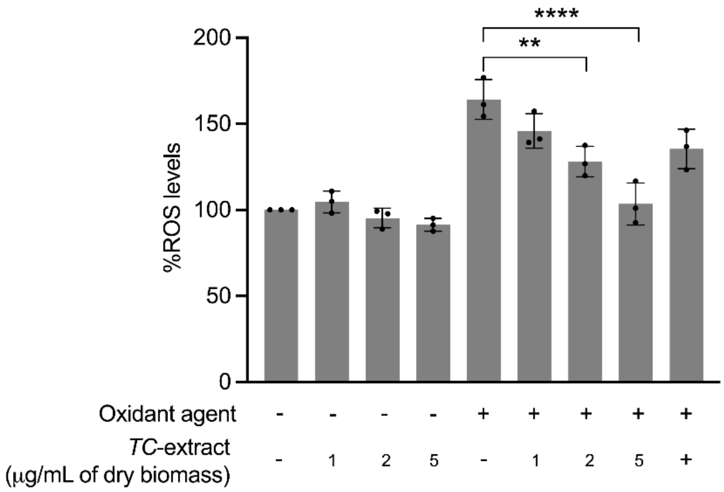


**Supplementary Figure 1: Antioxidant activity of *TC*-extracts.** The graph shows the percentage of intracellular ROS in NHDF cells, following treatment with *TC*-extract at 1, 2, 5 µg/mL of dry biomass, with or without the oxidative agent (250 µM TBH). *TC*-extract showed a strong antioxidant power, rebalancing ROS levels. The statistical analysis was performed by one-way ANOVA (oxidant agent *vs.* treatments), **$p < 0.01$, ****$p > 0.0001$.

### Supplementary Figure 2

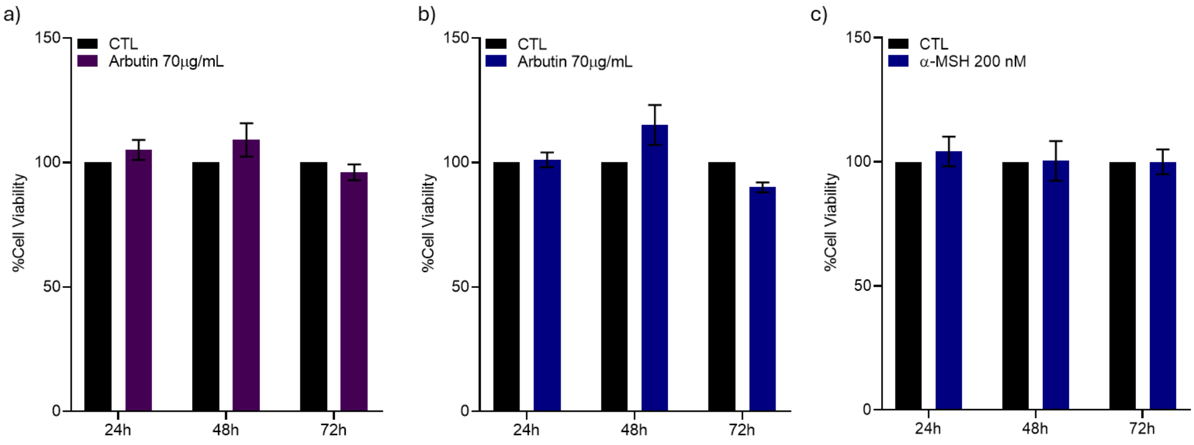


**Supplementary Figure 2: Biocompatibility assay of arbutin (70 µg/mL) and a-MSH (200 nM) in skin cells.** The graphs show the viability assay performed using MTS (3-(4,5-dimethylthiazol-2-yl)-5-(3-carboxymethoxyphenyl)-2-(4-sulfophenyl)-2H-tetrazolium) to: a) exclude any toxic effect of 70 µg/mL of arbutin treatment in NHDF and b) A375 cell lines, and c) to confirm the absence of toxic effect of 200 nM α-MSH treatment in A375. Cells were seed at a density of $2x10^3$ in 96-well plates and maintained in culture for 24h at 37°C 5% $CO_2$. Afterwards, cells were treated with arbutin and leave for MTS evaluation for 24, 48 and 72 hours. The statistical analysis was performed by two-way ANOVA (treatments *vs*. control).

Supplementary Figure 3

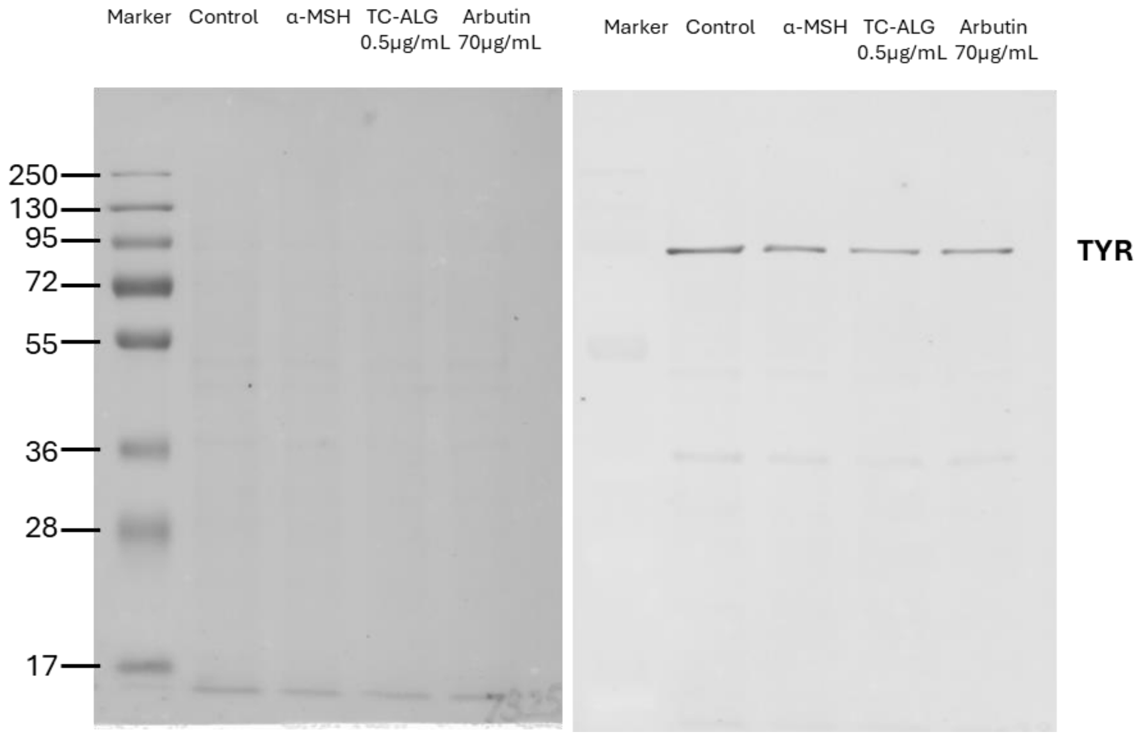


**Supplementary Figure 3: Unedited and uncropped western blot and corresponding Ponceau-stained blots.** The images show the western blot analyses for tyrosinase (TYR) expression evaluation after α-MSH stimulation.
